# The Largest Subunit of Human TFIIIC Complex, TFIIIC220, a Lysine Acetyltransferase Targets Histone H3K18

**DOI:** 10.1101/513127

**Authors:** Moumita Basu, Ramachandran Boopathi, Sadhan Das, Tapas K Kundu

**Author notes:** **Present Addresses:** Université Grenoble Alpes, CNRS UMR 5309, INSERM U1209, Institute for Advanced Biosciences (IAB), Site Santé – Allée des Alpes, 38700 La Tronche, France; Université de Lyon, Ecole Normale Supérieure de Lyon, CNRS, Laboratoire de Biologie et de Modélisation de la Cellule LBMC, 46 Allée d’Italie, 69007 Lyon, France. Department of Diabetes Complications and Metabolism, Diabetes Metabolism Research Institute, Beckman Research Institute of City of Hope, Duarte, CA 91010, USA. Correspondence: Prof. Tapas K Kundu, Transcription and Disease Laboratory, Molecular Biology and Genetics Unit, Jawaharlal Nehru Centre for Advanced Scientific Research, Bangalore 560064, India.

## Abstract

TFIIIC is a multisubunit complex that recognizes promoter elements and recruits TFIIIB and RNA polymerase III. Human TFIIIC complex possess lysine acetyltransferase activity which is critical in relieving chromatin mediated repression for RNA polymerase III-mediated transcription; two subunits of the TFIIIC complex, TFIIIC110 and TFIIIC90, were shown to acetylate H3 *in vitro.* Here we show that the largest and DNA binding subunit of TFIIIC complex, TFIIIC220, possesses intrinsic lysine acetyltransferase activity and acetylates histone H3K18 residue. By employing homology search we have identified the potential catalytic domain of TFIIIC220 which efficiently acetylate core histones *in vitro*. Point mutations at the critical residues of the identified acetyltransferase domain drastically reduces the acetyltransferase activity. Significantly, knockdown of TFIIIC220 in HepG2 cell line dramatically reduces global H3K18 acetylation level suggesting that TFIIIC220 is a crucial KAT to maintain acetylation homeostasis in the cell.

## INTRODUCTION

Acetylation of protein is one of the most prevalent posttranslational modification in eukaryotic system. Co-translational N_α_-acetylation occurs in almost ~80% of proteins which regulates their interaction, localization and stability (*Aksens et al., 2010*). Posttranslational addition of acetyl group from acetyl-CoA to N_ε_ of lysines is mostly catalyzed by lysine acetyltransferases (KATs) and has been classically associated with permissible chromatin structure, hence, transcriptional activation; but, later has also been shown to be critical for chromatin architecture (*Shogren-Knaak et al., 2006)*, DNA repair (*Chatterjee et al., 2012)*, protein stability and protein-protein interaction (*Kouzarides, 2007*). Currently, over 35,000 acetylation sites exist in human cells; abundance of this modification is almost comparable to that of protein phosphorylation (*Hornbeck et al., 2012*).

Eukaryotes possess very distinct set of nuclear lysine acetyltransferases which are majorly categorized based on structural and biochemical features of catalysis into the following subclasses; GCN5-related N-acetyltransferases (GNAT), the p300/CREB-binding protein (p300/CBP), and, the MOZ, Ybf2, Sas2, and Tip60 (MYST) family. However, some acetyltransferases which do not belong to any of these classes have also been identified later; among those there are nuclear hormone-related KATs SRC1 and ACTR (SRC3) and transcription factors such as human TFIID subunit TBP associated factor 250 (TAF_II_250) (*reviewed in Torchia et al., 1998*). General transcription factors like TFIIB and TFIIF has been found to possess autoacetylation properties which induces RNA polymerase II mediated transcription (*reviewed in Choi et al., 2004*). Also, most KATs belong to multiprotein complexes with various associated subunits, which regulate their catalytic activities and substrate specificities, especially in the case of histone acetylation (*Shahbazian et al., 2007*). Acetylation is a reversible process; metal-dependent lysine deacetylases (KDACs) catalyze the hydrolysis of acetyl-L-lysine side chains in proteins to yield L-lysine and acetate. Acetylation and deacetylation of nucleosomal histones provide a balance between open and closed chromatin conformation for the regulation of transcription. Like KATs, KDACs lack intrinsic DNA-binding activity and are recruited to target genes via their direct association with transcriptional activators and repressors, as well as their incorporation into large multiprotein transcriptional complexes (reviewed in *Lombardi et al., 2011*).

RNA polymerase III (pol III) transcribes small, untranslated structural RNAs contributing to ~15% of total RNA by weight. RNAPIII can initiate transcription from at least four different types of promoters; two of them being internal control region situating downstream of transcription start site (TSS) (*Dieci et al., 2007; Schramm et al., 2002*). Transcription from these two promoters essentially requires one general transcription factor, a multisubunit complex, TFIIIC. Once bound to promoter region TFIIIC can recruit TFIIIB and RNA polymerase III and form preinitiation complex. TFIIIC has been shown to be required for reinitiating transcription cycle and context dependent repression of RNA polymerase III mediated transcription. Other than RNA polymerase III mediated transcription TFIIIC has also been shown to possess barrier or insulator function, facilitate transposon insertion, localize in perinucleolar heterochromatin, facilitate or repress adjacent RNA polymerase II targeted transcription. TFIIIC has genome wide footprint majority of which is devoid of bound active RNA polymerase III, and, mediates ‘extra transcriptional effects’. TFIIIC may act as a stably bound, global “bookmark” within chromatin to establish, maintain, or demarcate chromatin states as cells divide or change gene expression patterns (*Policarpi et al., 2017; reviewed in Donze, 2011 and Van et al., 2012*). Human TFIIIC is ~600kDa multisubunit complex existing in different forms comprised of various subunits in cells (*Oettel et al., 1997*). Active TFIIIC comprises of at least six subunits (*Lagna et al., 1994; Ducrot et al., 2006*). The largest subunit is a 200kDa protein which recognizes and tightly binds to B box subunit through its N-terminal zinc finger binding motifs. Although functionally TFIIIC is conserved from yeast to human, TFIIIC220 homolog is not present in yeast; instead of 220kDa subunit a polypeptide of 138kDa binds to B-box region in yeast. Human TFIIIC holocomplex possesses acetyltransferse activity which aids in relieving chromatin mediated repression during transcription; TFIIIC can acetylate free and nucleosomal histones H3 and H4 in solution and can acetylate H2A as well in native HeLa nucleosome (*Kundu et al., 1999*). In this regard TFIIIC was found to be different from other general transcription factors possessing acetyltransferase activity such as TFIID subunit TAF_II_250 which can only acetylate H3 and H4 (*Mizzen et al., 1996*). Intrinsic acetyltransferase activities of two of TFIIIC subunits TFIIIC90 and TFIIIC110 have been characterized, their catalytic activities are specific for H3 (*Kundu et al.,1999; Hsieh et al.,1999*). However, if TFIIIC220 is an acetyltransferase or not is still unclear. A TFIIIC220 like protein in *C. tentans*, P2D10 has acetyltransferase activity specific for H3 and H4 *in vitro (Sjölinder et al., 2005*). This prompted us to investigate the acetyltransferase activity of human TFIIIC220. To understand whether TFIIIC220 is an acetyltransferase and its functional significance we purified recombinant human TFIIIC220 as well as its acetyltransferase domain and showed that hTFIIIC220 has intrinsic acetyltransferase activity. Most significantly, we showed that TFIIIC220 is an acetyltransferase with strong specificity for H3K18 residue *in vitro* as well as in the cellular context.

## RESULTS AND DISCUSSION

### TFIIIC220 has intrinsic acetyltransferase activity

In-gel histone acetyltransferase assay indicated that the human TFIIIC complex possesses three polypeptides, 220, 110 and 90kDa with intrinsic acetyltransferase enzymatic activity. The catalytic activities of TFIIIC110 and TFIIIC90 were confirmed using recombinant deletion mutation domains or baculovirus expressed full length protein and were shown to efficiently acetylates core histones (*Kundu et al., 1999; Hseih et al., 1999*) However, the catalytic activity of the largest subunit of TFIIIC complex, TFIIIC220 is yet to be established. Since TFIIIC complex was found to pull down robust acetyltransferase p300 (*Mertens et al., 2008*) it might also be the factor contributing to the acetyltransferase activity of the complex. To characterize the contribution of catalytic activity of the 220 kDa subunit we constructed the recombinant baculovirus using full length clone of human TFIIIC220and purified it from Sf21 cells infected with the baculovirus containing His-tagged full length human TFIIIC220 isoform 1 (Figure 1A). The expression of the protein was confirmed by TFIIIC220 specific antibody (Figure 1A). Immunoblottting the eluted fraction with p300 specific antibody confirmed that eluted fraction of TFIIIC220 was not contaminated with p300 from Sf cells (Supplementary figure 1). We next examined intrinsic catalytic activity of TFIIIC220 by *in vitro* enzymatic assays using recombinant core histones H3, H2A and H4, since substrate specificity of TFIIIC holocomplex were found to be specific for histones H3, H4 and H2A but not for H2B. Full length TFIIIC220 was found to possess significant amount of enzymatic activity (Figure 1B) and it could acetylate recombinant H3, H2A and H4 (Figure 1C).

**Figure 1:**
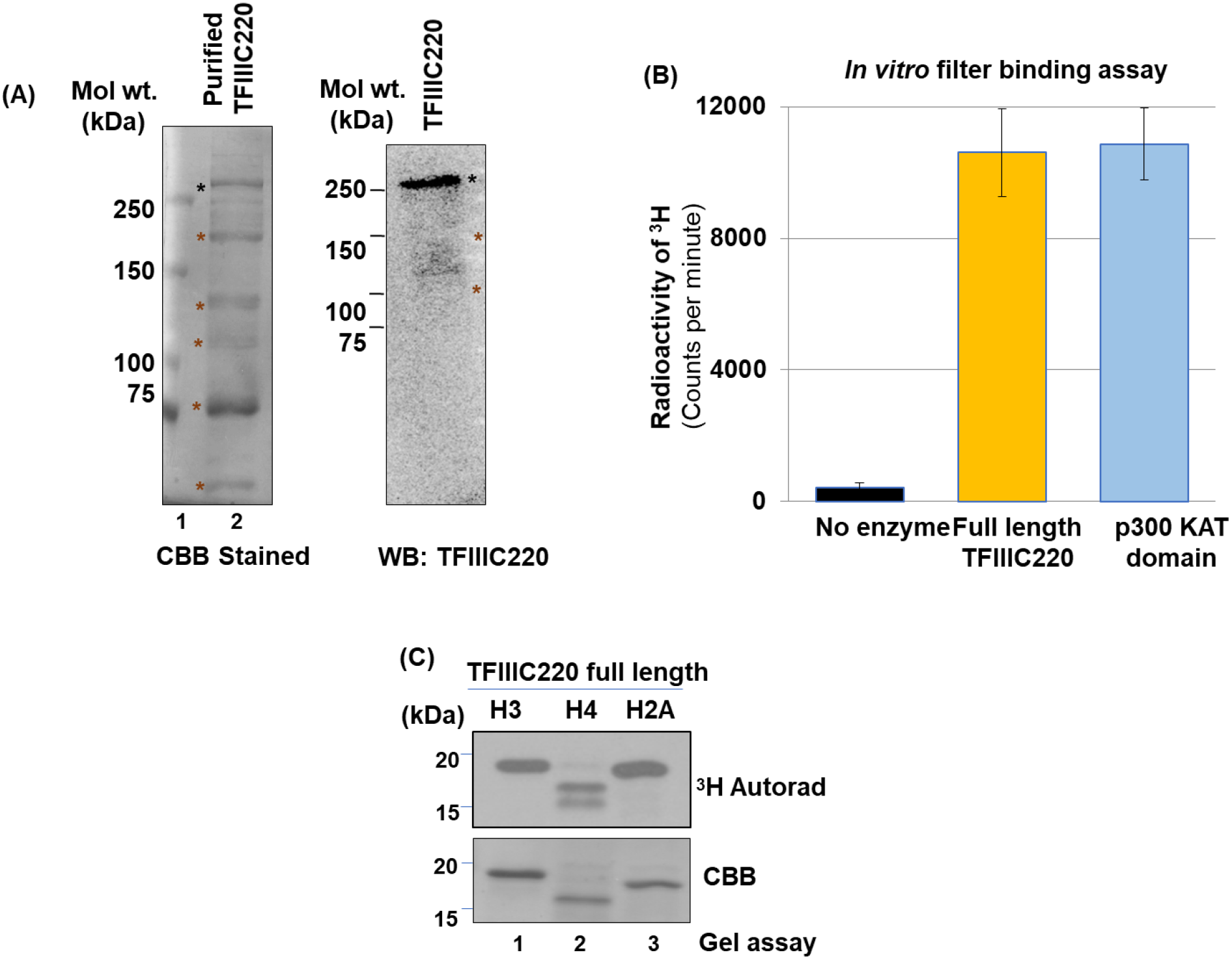
Recombinant TFIIIC220 possesses acetyltransferase activity. **A.** SDS PAGE profile of purified full length human TFIIIC220. Purified protein was probed with anti-His polyclonal antibody. TFIIIC220 specific antibody recognizes the intact protein of ~250kda (indicated with black star) as well as some of the degraded products (indicated with brown stars). **B.** *In vitro* filter binding assay was performed using full length TFIIIC220 and human p300 KAT domain. 1μg histone H3 and 50nCi ^3^H-acetyl CoA were used as substrates. **C.** *In vitro* KAT assay was performed using full length TFIIIC220, 1μg of histone H3, H4 and H2A and 50nCi ^3^H-acetyl CoA were used as substrates. Reaction mixtures were loaded onto 12% SDS-PAGE and developed on X ray films.

### TFIIIC220 KAT activity maps to predicted acetyl CoA binding sites

We next mapped TFIIIC220 domains responsible for its acetyltransferase activity. TFIIIC220 did not show any sequence similarity with known acetyltransferases, but three regions of human TFIIIC220 matched the highly conserved acetyl CoA recognition and binding motif (R/Q)XXGX(G/A) (*Dutnall et al., 1998*); Two of those aa 834-839 and aa 896-901 lies in the mid region and motif 3 aa 1941-1946 is in the C terminal of isoform 1 (Figure 2B); Interestingly, we found that isoform 2 of human TFIIIC220 lacks putative motif 3. Motif 1 and 2 are 100% conserved among mammals but unlike motif 1, motif 2 is conserved across other vertebrates as well (Supplementary figure 2 and Figure 2B). We considered ~50kDa region spanning motif 1 and 2 to be the putative acetyltransferase domain and purified recombinant KAT domain from *E.coli (Figure 2C*). Purified protein was identified by immunoblotting with antibodies specific for His tag which recognized the protein corresponding to ~50kDa (Figure 2C). 50kDa protein was also confirmed to be the putative acetyltransferase domain of hTFIIIC220 by mass spectrometric analysis (Supplementary figure 3). Additional proteins in the elution corresponding to molecular weights of ~25 and ~75kDa were identified as common histidine rich contaminants from BL21 cells by mass spectrometric analysis (Data not shown, as mentioned in Bolanos-Garcia et al., 2006). Purified TFIIIC220 KAT domain possessed significant acetyltransferase activity and acetylated histone H3 *in vitro* (Figure 2D). Next, we introduced point mutations in putative acetyl CoA binding motif to destabilize interaction with acetyl CoA by mutating glycines with electronegative and bulky aspartic acid and arginines with small neutral amino acid alanine. Introduction of destabilizing point mutations in putative acetyl CoA binding motif 1 abrogated its acetyltransferase activity by ~40% in *in vitro* enzymatic assay (Figure 2D). The remnant catalytic function might result from intact motif 1 present in the putative catalytic domain.

**Figure 2:**
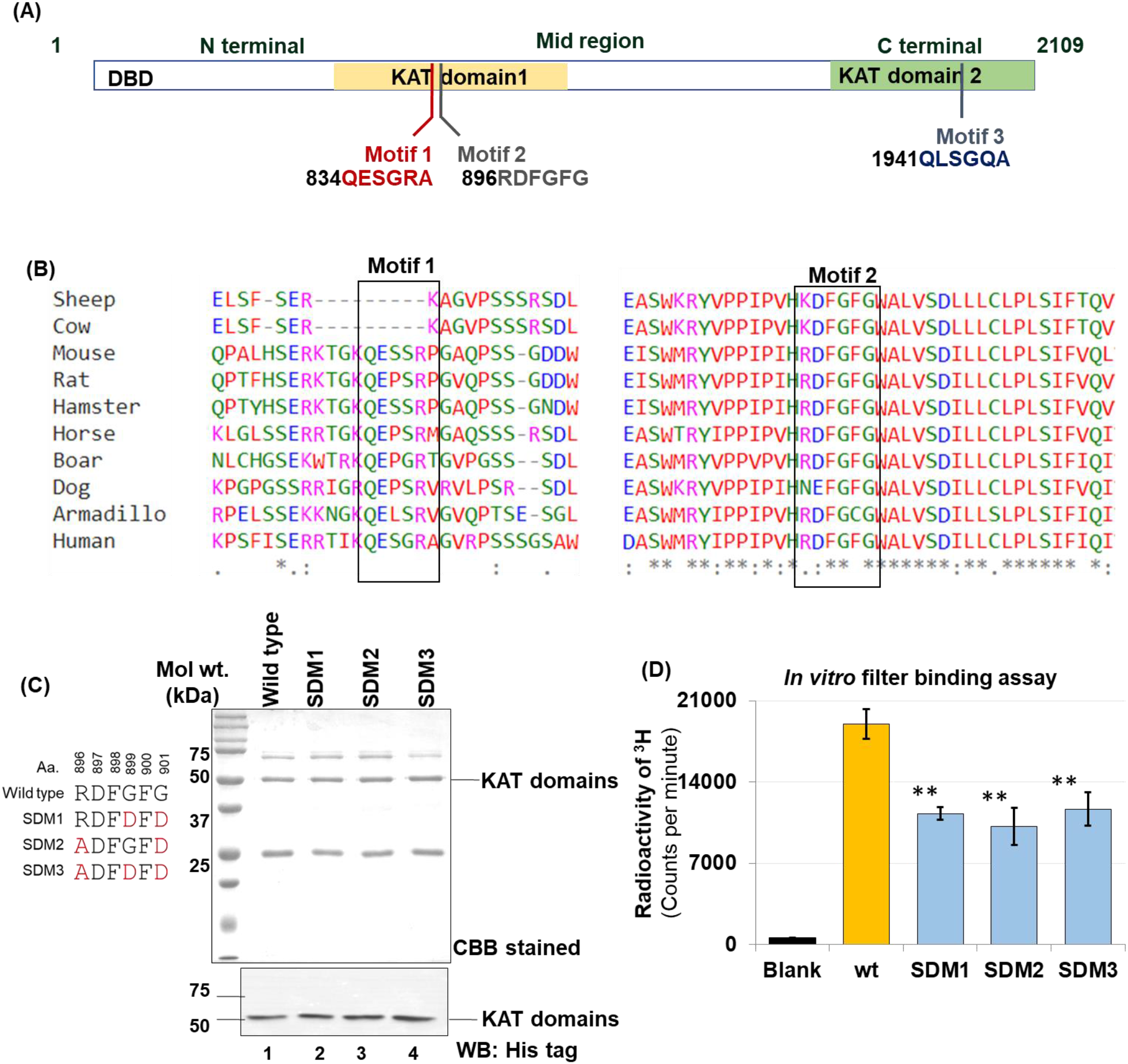
Identification of acetyltransferase domain of TFIIIC220. **A.** Schematic diagram of full length TFIIIC220 protein. Relative positions of three putative acetyl CoA binding motifs are indicated. [DBD= DNA binding region] **B.** Sequence conservation of motif 1 and motif 2 across different vertebrate species using Clustal Omega (EMBL-EBI). **C.** SDS-PAGE profile of purified recombinant KAT domain and mutants. D. *In vitro* filter binding assay was performed using ~200 ng wild type and mutant TFIIIC220 KAT domains, 1μg of histone H3, H4 and H2A and 50nCi ^3^H-acetyl CoA were used as substrates. Student’s unpaired t test was used for statistical analysis; N=2; p ns>0.05, p*<0.05, p**<0.001, p***<0.0005

**Figure 3:**
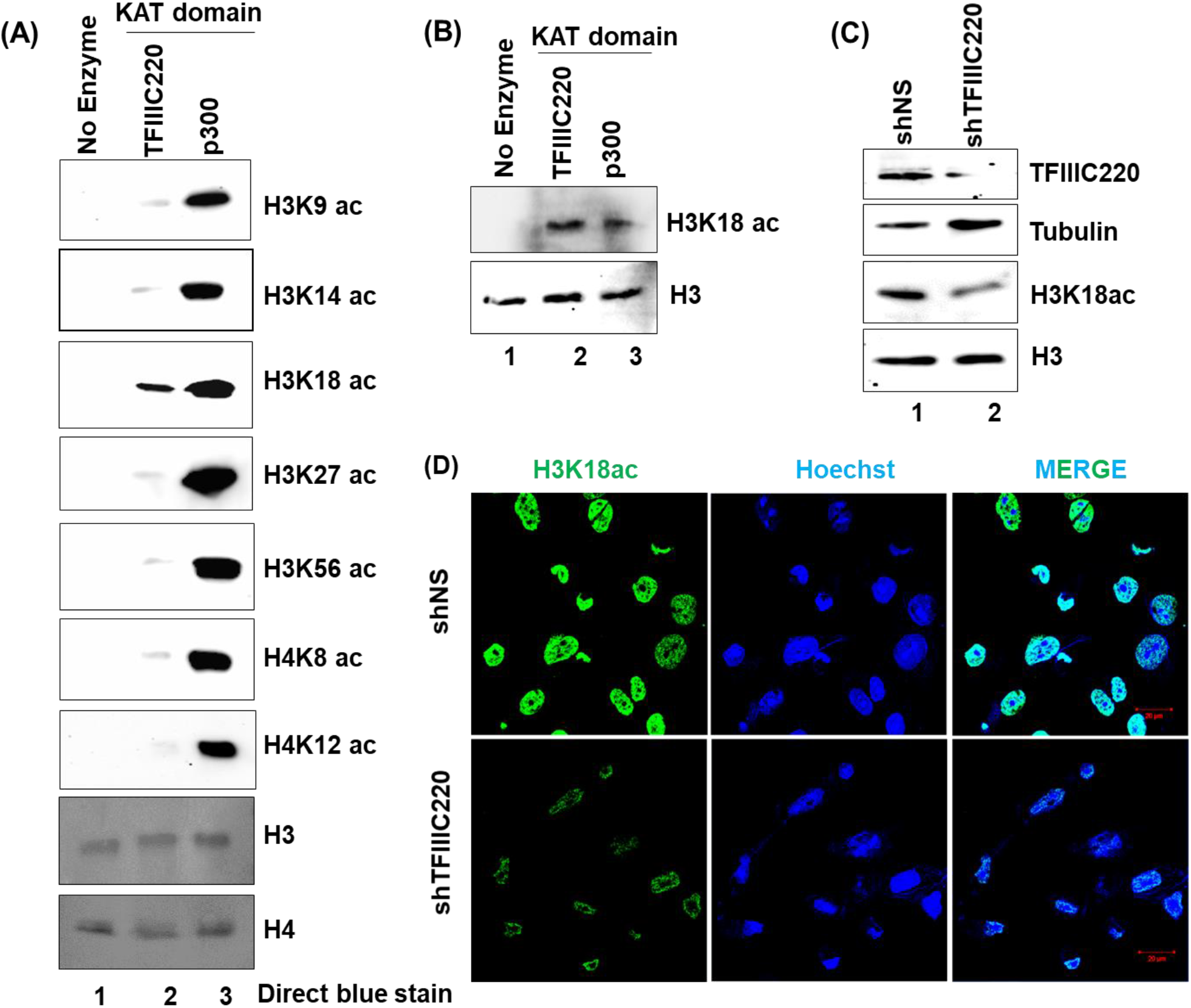
Site specificity of TFIIIC220 on core histones *in vitro*. ***A.** In vitro* KAT assay was performed using ~200ng of recombinant TFIIIC220 KAT domain. ~1μg of recombinant H3, H4 and 40μM acetyl CoA were used as substrates. Reaction mixtures were loaded on 12% SDS PAGE and probed with site specific acetyllysine antibodies. Human p300 KAT domain was taken as positive control for all the reactions. **B.** *In vitro* KAT assay was performed using reconstituted nucleosome and then loaded on 12% SDS PAGE and probed with acetylated H3K18 specific antibody. Human p300 KAT domain was taken as positive control. **C and D**. HepG2 cells were stably transfected with shTFIIIC220 or shNS (scrambled) and treated with 2μg/mL doxycycline for 96 hours. RIPA and Laemmli lysates were loaded onto 12% SDS PAGE and probed with TFIIIC220 (normalized against tubulin) and acetylated H3K18 specific antibodies (normalized against histone H3) (in C); cells were probed with acetylated H3K18 specific antibody and observed under microscope for immunofluorescence. Scale bar: 20μm.

### TFIIIC220 KAT differs from known acetyltransferases

Effects of acetyltransferase activities of different KAT families have been found to be diverse. Although in general acetylation of histones opens up the chromatin structure and activates transcription, KATs have also been shown to have implication in DNA repair, protein stabilization, protein-protein interaction *etc*. Since, KAT domain of TFIIIC220 did not bear similarities with other known classes of nuclear KATs we expected its acetylation signature to be different as well. In order to characterize the residue specificity of the KAT activity of TFIIIC220 recombinant acetyltransferase domain of TFIIIC220 was used for *in vitro* acetyltransferase assay. Since p300 is one the most robust and well conserved acetyltransferase known to acetylate all the core histones in many lysine residues we used recombinant human p300 KAT domain as positive control for our assays. The results showed that TFIIIC220 KAT domain could efficiently acetylate H3K18 residue which is known to be associated with open chromatin and active transcription. Surprisingly TFIIIC220 KAT domain failed to modify other N -terminal lysine residues of H3 such as K9, K14, K27, and, K56 and H4 K8 and K12 *in vitro* (Figure 3A). Since specificity of KATs for histones could be close to the *in vivo* context when nucleosomes are used as substrates we reconstituted mononucleosomes using HeLa cells core histones and used as substrates in enzymatic assay. Full length TFIIIC220 also showed specific acetyltransferase activity towards reconstituted mononucleosome and acetylated at H3K18 residue. Recombinant full length TFIIIC220 could not acetylate H3K14 and H3K56 residues *in vitro* further indicating it is free from p300 contamination (Supplementary figure 4). To address the *in vivo* specificity of KAT activity of TFIIIC220 we downregulated TFIIIC220 expression in HepG2 cells by TFIIIC220 specific shRNA stable transfection and probed for H3K18 acetylation in TFIIIC220 knockdown cells. Knockdown of TFIIIC220 drastically reduced global H3K18 acetylation level in HepG2 cells (Figure 3C and D).

**Figure 4:**
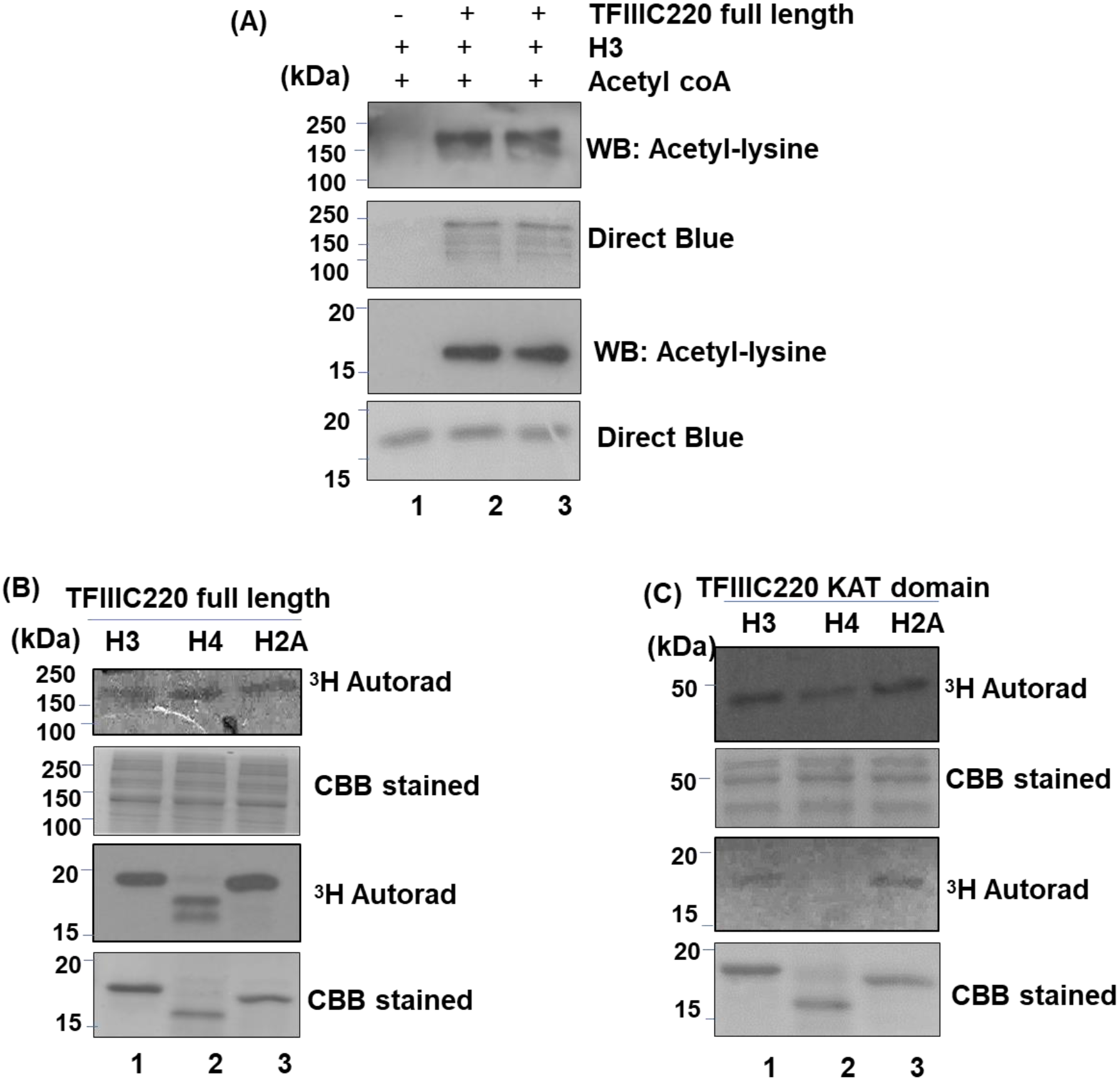
TFIIIC220 autoacetylates. *In vitro* KAT assay was performed using ~100 ng full length and ~200ng KAT domain of TFIIIC220. 1μg of recombinant core histones (H3, H2A, and, H4) and 40μM acetyl CoA or 50nCi ^3^H-acetyl CoA were used as substrates. Reaction mixtures were loaded onto 12% SDS PAGE and probed with acetyl-lysine specific antibody (in A) or developed onto X ray films (in B and C)

### TFIIIC220 has autoacetylation property

Like kinases the lysine acetyltransferases also possess self-enzymatic activity. Most of the KATs have been shown to be autoacetylated which enhance their catalytic activity. We therefore investigated whether TFIIIC220 possesses autocatalytic property. We found that purified full length TFIIIC220 from Sf cells are highly acetylated or undergoes rapid acetylation under *in vitro* assay conditions (Figure 4A). To distinguish the cellular acetyl CoA pool from that of the assay we used tritiated acetyl CoA and found that full length TFIIIC220 as well as its KAT domain stably incorporated tritiated acetyl group in its structure (Figure 4B and C).

This study has provided experimental evidence to establish TFIIIC220 as a bona fide histone acetyltransferase. Full length TFIIIC220 could acetylate nucleosomal and free histone H3 *in vitro* and specifically acetylated H3K18 residue. Earlier TFIIB and TFIIF, general transcription factors for RNA polymerase II was found to possess acetyltransferase activity, they regulated themselves by autoacetylation and their catalytic activities have implication in transcription (*Choi et al., 2004*). Our study will be a first finding of a general transcription factor which directly binds to DNA also possesses histone acetyltransferase activity along with autoacetylation property. Given the versatility of TFIIIC complex intrinsic catalytic activities of its subunits might play important roles in orchestrating its multitudinal effector function. TFIIIC220 being the DNA binding subunit and main recruitment factor of TFIIIC holocomplex we presume that acetyltransferase activity of this subunit holds special significance. Acetylation of different lysine residues of histones is closely linked to various biological phenomena. TFIIIC220 preferentially acetylates Lys18 in H3 N-terminal. Lys18 in H3 is also a preferred substrate for acetylation by human p300/CBP and ELP3. Acetylation of this particular site is shown to be enriched in active TSS, implicated in nucleolar heterochromatin dynamics, cytotoxicity and poor prognosis in cancer (*Juliano et al., 2016; Damodaran et al., 2017; Hiraoka et al., 2013; Ianni et al., 2017*)

TFIIIC220 is a higher eukaryote protein, no homologue of TFIIIC220 is found in yeast. Although a functional homolog of TFIIIC220 is identified in *Drosophila* it does not bear significant sequence similarity and acetyltransferase activity of the protein is not characterized. TFIIIC220 in vertebrates are conserved more or less; the homologs being almost identical in mammals gives rise to the possibility of highly evolved function of the protein.

## MATERIALS AND METHODS

### Cell lines and culture

HepG2 cells (ATCC) were grown in MEM (Gibco) with 2mM Glutamine, 10% FBS at 37 °C and 5.0% CO2. Cell lines were tested for mycoplasma contamination. Sf21 cells (Invitrogen) were grown at 27 °C in Grace’s Insect Medium (Gibco) with 10% FBS.

### Stable cell line generation

shTFIIIC220 or scrambled shRNA construct (Dharmacon) containing lentivirus were generated using calcium phosphate mediated precipitation of 5μg of pSPAX2, 0.75μg pCMV-Rev, 1.75μg pVSVG and 5μg of respective shRNAs in HEPES buffered saline (pH 7.05) onto HEK293T cells in 90mm dish containing 10mL of DMEM supplemented with 10% FBS. Stable cell line was generated by infecting HepG2 cells with lentivirus containing shRNAs for ~8 hours. Cells were selected with 1 mg/mL Puromycin (Sigma) and expression of shRNAs were monitored under microscope by RFP expression after 96 hours treatment with 2mg/mL Doxycycline (Sigma).

### Plasmid constructs

PTRF-IIICa clone was a kind gift from Dr. Zhengxin Wang, MD Anderson Cancer Research Centre, Texas. TFIIIC220 full length construct was subcloned into pENTR-DTOPO vector using Topo TA cloning kit (Invitogen) and primers listed below. TFIIICFP topo 5’ CACCATGGACGCGCTGGAGTC 3’, Topo 1^st^ RP 5’ CTCGAGCGGAATTCCGCTCTAGAGGAAGCACTCAGCT 3’, Topo 2^nd^ FP 5’ C TAGTCTAGATCAGAAAGT GGACGGATGAAAAAAAG 3’, Topo 2^nd^ RP: 5’CGGAATTCCTTAGGGCTGAACTGAACTTTTC 3’, Topo 3^rd^ FP: 5’ CGGAATTCTAACCTTGAAATCCCAGACACAC 3’, Topo 3^rd^ RP: 5’ CCGCTCGAGGAGGTGGATCCACTTG 3’, TFIIICRP topo 5’ GAGGTGGATCCACTTGTTCCAGTTGACC 3’. Clone was confirmed by sequencing and cloned into baculovirus using BaculoDirect Bacoluvirus Expression system kit (Invitrogen). C-terminally His6 tagged TFIIIC220 KAT domain was cloned in the pET28b vector using PCR based cloning from TFIIIC220 full length clone. Catalytic mutants were generated by site directed mutagenesis (Agilent Quik Change II Mutagenesis Kit). Primers used: GTF3C1 KAT domain forward: 5’ CATGCCATGGCAATGTTTCTGTGGTAC 3’, GTF3C1 KAT domain reverse: 5’ CCGCTCGAGAACCAGTAGTTTTCCAC 3’, SDM1 forward: 5’ GTGTCCCTGAAGCTGAAACTGACCCGAGAGCAG 3’, SDM1 reverse: 5’ CACAGGGACTTCGACTTTGACTGGGCTCTCGTC 3’, SDM2 forward: 5’ GAGCCCAGTCAAAGTCGAAGTCCGCGTGG 3’, SDM2 reverse: 5’ CCACGCGGACTTCGACTTTGACTGGGCTC 3’, SDM3 forward: 5’ GAGCCCAGTCAAAGTCGAAGTCCGCGTGG 3’, SDM3 reverse: 5’ CCACGCGGACTTCGACTTTGACTGGGCTC 3’

### Recombinant protein expression and purification

Protein purification protocol was adapted from *Bornhorst and Falke, 2000*. Full length TFIIIC220 was purified from Sf21 cells infected with TFIIIC220 baculovirus for 72 hours. Recombinant wildtype and mutant KAT domains of TFIIIC220 were co-expressed with Sirt2 in *E. coli* BL21 (DE3) cells to reduce toxic effect of KAT overexpression and induced with 0.4 mM IPTG at 30 °C for 5 h. Cells were lysed in BC300(20 mM Tris-Cl pH 7.4, 20% Glycerol, 300 mM KCl, 0.1% NP40, 2 mM PMSF, 2 mM beta mercaptoethanol, Protease Inhibitor Cocktail (Sigma)) and sonicated until lysate was clear. Lysates were incubated with Ni-NTA beads (Novagen) for 3 h and washed with BC300. Proteins were eluted in BC100 containing 250 mM Imidazole. 6His-tagged p300 HAT domain (aa 1284-1673) was purified from cells co-transformed with p300 HAT domain and SirT2 construct in E. coliBL21(DE3) as described previously in *Thompson et al., 2004*.

### Reagents

Antibodies used for immunoblots and immunofluorescence are anti-TFIIIC220 (Bethyl Laboratories, A301-291A), anti-tubulin (Calbiochem, CP06), histone modifications antibodies were raised in-house. Secondary antibodies: Goat Anti-Rabbit IgG H&L (HRP), Abcam (Catalog no. ab97051); Goat anti-Rabbit IgG (H+L) Cross-Adsorbed, Alexa Fluor^®^ 488, Thermo Fisher Scientific (Catalog no. A-11008).

### *In vitro* filter binding assay and gel assay

Filter binding HAT assay was performed using 50nCi 3H acetyl-CoA and 1μg recombinant H3 or ~1μg of reconstituted nucleosome as substrate (*Berndsen and Denu, 2005*) and analyzed by capturing the acetylated products on Whatman p81filter paper. After stringent washes to remove residual free acetyl CoA, acetylation levels were quantified with Wallac 1409 Liquid Scintillation Counter. For gel assay reaction mixtures were loaded onto 12% SDS PAGE and then transferred onto PVDF membrane (Merck Millipore) and immunoblotted with antibodies as mentioned. Nucleosome reconstitutions were performed by salt gradient dialysis as described previously (Hayes et al., 1997) 601 DNA (kind gift from Jonathan Widom) amplified from pGEM3z clone using appropriate primers (listed below) and HeLa core histones and free DNA in 1:1 molar ratio were incubated in Buffer A (10mM Tris-HCl pH 7.9, 1mM EDTA, 1mM β-Me) containing 2M NaCl. Reaction was step dialysed in Buffer A containing 1, 0.8, 0.6, 0.1 M NaCl. Finally reaction was dialysed against Buffer B (10mM Tris-HCl pH 7.9, 0.25mM EDTA, 10mM NaCl) for overnight at 4ºC. 601 FP: 5’ GCTCGGAATTCTATCCGACTGGCACCGGCAAG 3’ 601 RP: 5’ GCATGATTCTTAAGACCGAGTTCATCCCTTATGTG 3’

## Acknowledgement

We acknowledge JNCASR confocal facility for confocal imaging and IISc Bangalore MBU mass spectrometric facility for LC-MS service.

