## Supplementary Figures for "The Largest Subunit of Human TFIIIC Complex, TFIIIC220, a Lysine Acetyltransferase Targets Histone H3K18"

**Supplementary Information**

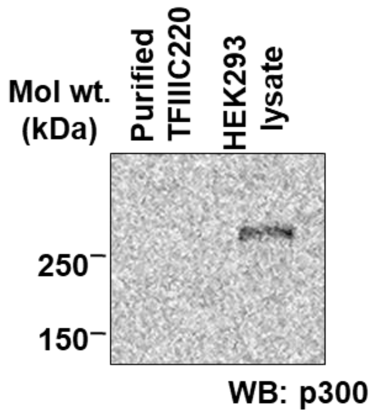


**S.Fig.1: Purified fraction of full length TFIIIC220 was probed with p300 specific antibody (Santacruz, sc-585); HEK293 cell lysate was used as positive control for p300 immunoblotting.**

| Class | Vertebrate Organism (Ortholog name: TF3C1) | Sequence Identity (%) | Motif 1 (QESGRA) | Motif 2 (RDFGFG) | Motif 3 (QLSGQA) |
|---|---|---|---|---|---|
| Fish | Zebrafish (511aa) | 36.38 | | | |
| Amphibian | Xenopus tropicalis (1999aa) | 56.27 | | | |
| Reptile | Lizard (2131aa) | 61.56 | | | |
| Bird | Fowl (2145aa) | 66.23 | | | |
| | Mallard (2099aa) | 67 | | | |
| Mammal | Sheep | 78.21 | ----KA | KDFGFG | RVGPTS |
| | Hamster | 78.56 | QESSRP | RDFGFG | QLGCEF |
| | Rat | 79.17 | QEPSRP | RDFGFG | QLGCEF |
| | Mouse | 79.25 | QESSRP | RDFGFG | QLGCEF |
|  | Cow | 80.21 | ----KA | KDFGFG | QLSCQA |
|  | Pig | 81.48 | QEPGRT | RDFGFG | QL---- |
| | Dog | 81.78 | QEPSRV | RDFGFG | QLSCQD |
| | Horse | 84.11 | QEPSRM | RDFGFG | QLSCQA |
| | Armadillo | 84.52 | QELSRV | RDFGCG | QPSCQA |
| | Green Velvet Monkey | 97.44 | QESGRA | RDFGFG | QLSGQA |
| | Rhesus macaque | 97.58 | QESGRA | RDFGFG | QLSGQA |
| | Olive baboon | 97.58 | QESGRA | RDFGFG | QLSGQA |
| | Sumatran Orangutan | 98.62 | QESGRA | RDFGFG | QLSGQA |
| | Gorilla | 99.07 | QESGRA | RDFGFG | QLSGQA |
| | Chimpanzee | 99.67 | QESGRA | RDFGFG | QLSGQA |
| | Human | 100 | QESGRA | RDFGFG | QLSGQA |

**S.Fig.2: Conservation of putative acetyl CoA binding motif across different species. Sequences of most prevalent TFIIIC220 isoforms or predicted homologs were collected from NCBI database and motifs were identified by aligning with hTFIIIC220 using Clustal omega software (EMBL-EBI)**

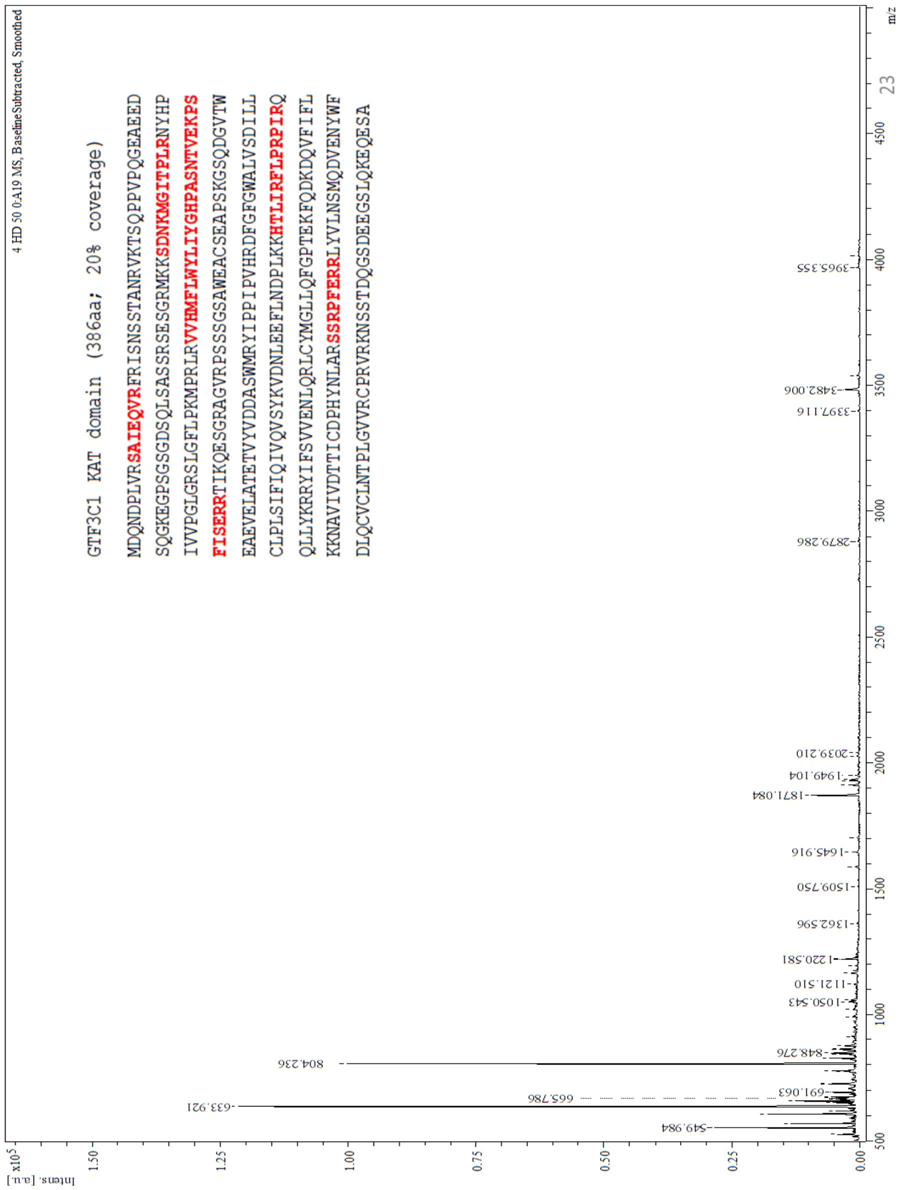


**S.Fig.3: Identification of 50kDa protein band in recombinant hTFIIIC220 KAT domain elution by LC-MS analysis. Aminoacid sequence of KAT domain is given in the subset and peaks corresponding to the tryptic digested fragments were indicated in red.**

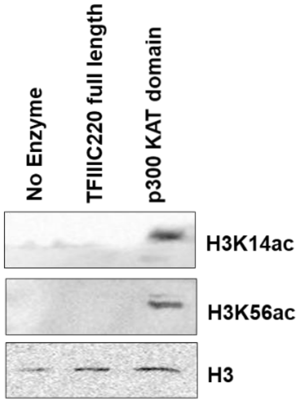


**S.Fig.4: Gel assay with reconstituted nucleosomes with HeLa core histones and 200ng full length TFIIIC220. p300 KAT domain was used as positive control for the assay. Reaction mixtures were loaded onto 12% SDS PAGE and probed with acetylated H3K14, K56 and H3 specific antibodies.**
